# Unravelling Different Biological Roles Of Plant Synaptotagmins

**DOI:** 10.1101/2024.01.21.576508

**Authors:** Selene García-Hernández, Lourdes Rubio, Jessica Pérez-Sancho, Alicia Esteban del Valle, Francisco Benítez-Fuente, Carmen R. Beuzón, Alberto P. Macho, Noemi Ruiz-López, Armando Albert, Miguel A. Botella

## Abstract

Endoplasmic Reticulum-Plasma Membrane contact sites (ER-PM CS) are evolutionarily conserved membrane domains found in all eukaryotes, where the endoplasmic reticulum (ER) closely interfaces with the plasma membrane (PM). This short distance, typically 10-30 nm, is achieved in Arabidopsis through the action of tether proteins such as Synaptotagmins (SYTs). Arabidopsis comprises five SYT members (*SYT1-SYT5*), but whether they possess overlapping or distinct biological functions remains elusive. *SYT1* is the best-characterized gene member and plays an essential role in the resistance to abiotic stress. This study reveals that while the functionally redundant *SYT1* and *SYT3* genes are involved in salt and cold stress resistance, *SYT5* is associated with *Pseudomonas syringae* resistance despite evidence of *in vivo* interaction between SYT1 and SYT5. Structural phylogenetic analysis indicates that SYT1 and SYT5 clades emerged early in the evolution of land plants. These protein clades exhibit different structural features, rationalizing their distinct biological roles.

## INTRODUCTION

Membrane contact sites (MCS) are microdomains where the membranes of two organelles closely approach each other, typically within the range of 10–30 nm (Fernández-Busnadiego et al., 2015). This proximity is achieved through proteins acting as molecular tethers, bridging the two membranes (Eisenberg-Bord et al., 2021; Pérez-Sancho et al., 2016). MCS are widespread in nearly all organelles, mostly involving the Endoplasmic Reticulum (ER). The minimal distance between membranes facilitates effective organelle communication, often leading to the exchange of molecules such as ions and lipids (Pérez-Sancho et al., 2016).

Plant synaptotagmins (SYTs), along with their ortholog counterparts, the mammalian extended synaptotagmins (E-Syts) and yeast tricalbins (Tcbs), serve as tethers and lipid transfer proteins between the ER and the plasma membrane (PM). These proteins share a common modular structure, including an N-terminal anchoring them to the ER, a SYT-like mitochondrial-lipid binding (SMP) domain, and a variable number of Ca^2+^ and phospholipid-binding C2 domains. SMP domains belong to the tubular lipid-binding protein superfamily, characterized by a folding structure that accommodates lipids in a hydrophobic cavity (Kopec et al., 2010). The shared modular structure of these proteins suggests common functions, including two closely related roles: the establishment of Ca^2+^-regulated ER-PM tethering facilitated by their C2 domains (Giordano et al. 2013; Manford et al. 2012; Pérez-Sancho et al. 2015; Saheki et al. 2016), and the SMP-dependent transport of lipids between the PM and the ER (Ruiz-Lopez et al., 2021; Saheki et al., 2016; Schauder et al., 2014)

The Arabidopsis SYT family consists of five members (*SYT1*-*SYT5). SYT1* is the best-characterized gene family member, and a loss-of-function mutant in *SYT1* shows multiple defects in abiotic stress responses, including salt (Schapire et al. 2008), freezing (Ruiz-Lopez et al., 2021; Yamazaki et al., 2008), and wounding (Pérez-Sancho et al., 2015), but also displays altered resistance to virus (Levy et al., 2015; Lewis & Lazarowitz, 2010; Uchiyama et al., 2014) and fungi (Y. J. Kim et al., 2016). Interestingly, while *SYT3* has been shown to have a redundant role with *SYT1* in abiotic stress, *SYT5* has been shown to have a role in the resistance to *Pseudomonas syringae* pv tomato (S. Kim et al., 2021a, 2021b). In addition, various mutant combinations cause slight defects in growth and seed production (Benitez-Fuente & Botella, 2023; Ishikawa et al., 2020). Because of their structural characteristics, the variable function of SYTs may depend on their tethering role, as shown by their role in ER stability (Siao et al., 2016). Still, it could also be caused by a role in lipid transport mediated by their SMPs. Regarding this, SYT1 and SYT3 proteins have been proposed to have a specific role in maintaining diacylglycerol homeostasis at the PM after the stress-activation of phospholipase C (Ruiz-Lopez et al., 2021).

Critical information regarding the role of plant SYTs involves determining whether they play overlapping or diverse biological functions. Surprisingly, this remains an open question not only in plant SYTs but also in the well-characterized mammals E-Syts and yeast Tcbs. Tether properties based on their C2 domains, specificities of the lipids transported by their SMPs, or different interactomes can account for specific biological roles. Here, we show that Arabidopsis *syt1* and *syt3* mutants show different phenotypes than *syt5*, supporting different biological roles within this gene family. Further phylogenetic and structural analyses support the divergence of *SYT1* and *SYT5* genes during the early stages of land plant evolution, explaining their different functionalities.

## RESULTS AND DISCUSSION

### Arabidopsis SYT1 and SYT5 play different roles in stress tolerance despite being part of the same complexes at ER-PM CS

The Arabidopsis genome contains five Synaptotagmin (*SYT*) genes, yet it is unknown whether all family members show similar biological functions. Expression analyses of *SYT* genes using available RNA sequencing data revealed that *SYT1* transcripts were approximately 4 and 17 times more abundant than *SYT5* and *SYT3*, respectively (Fig. S1A), whereas SYT4 and SYT2 exhibited low expression in vegetative tissues (Fig. S1B). Therefore, we focused on *SYT1/SYT3* and *SYT5* for further studies.

*SYT1* and *SYT3* play important and redundant roles in the resistance of Arabidopsis to various abiotic stresses (Benitez-Fuente & Botella, 2023; Ruiz-Lopez et al., 2021). Using fluorescein diacetate (FDA) as a vital stain (Schapire et al. 2008) we showed that 30 min exposure to cold treatment (8°C) did not cause an impact on the cell survival of WT, *syt3,* or *syt5* roots while induced a substantial reduction in cell viability in *syt1* roots (Fig. 1A). The highly expressed SYT1 could mask the possible role of *SYT5* in cell viability as occurred for *SYT3* (Ruiz-Lopez et al., 2021). Therefore, we generated a *syt1syt5* double mutant and performed a shorter cold treatment in which the double *syt1syt3* mutant showed enhanced cell death compared to *syt1* (Ruiz-Lopez et al., 2021). For this treatment, *t*en minutes of cold caused a mortality of around 80% in the *syt1syt3* double mutant, while *syt1* showed around 50% dead cells. However, the cell viability of the *syt1syt5* double mutant resembled that of *syt1* (Fig. 1B). This data indicates the absence of redundant function between *SYT1* and *SYT5* in cell viability under cold stress.

**Figure 1.**
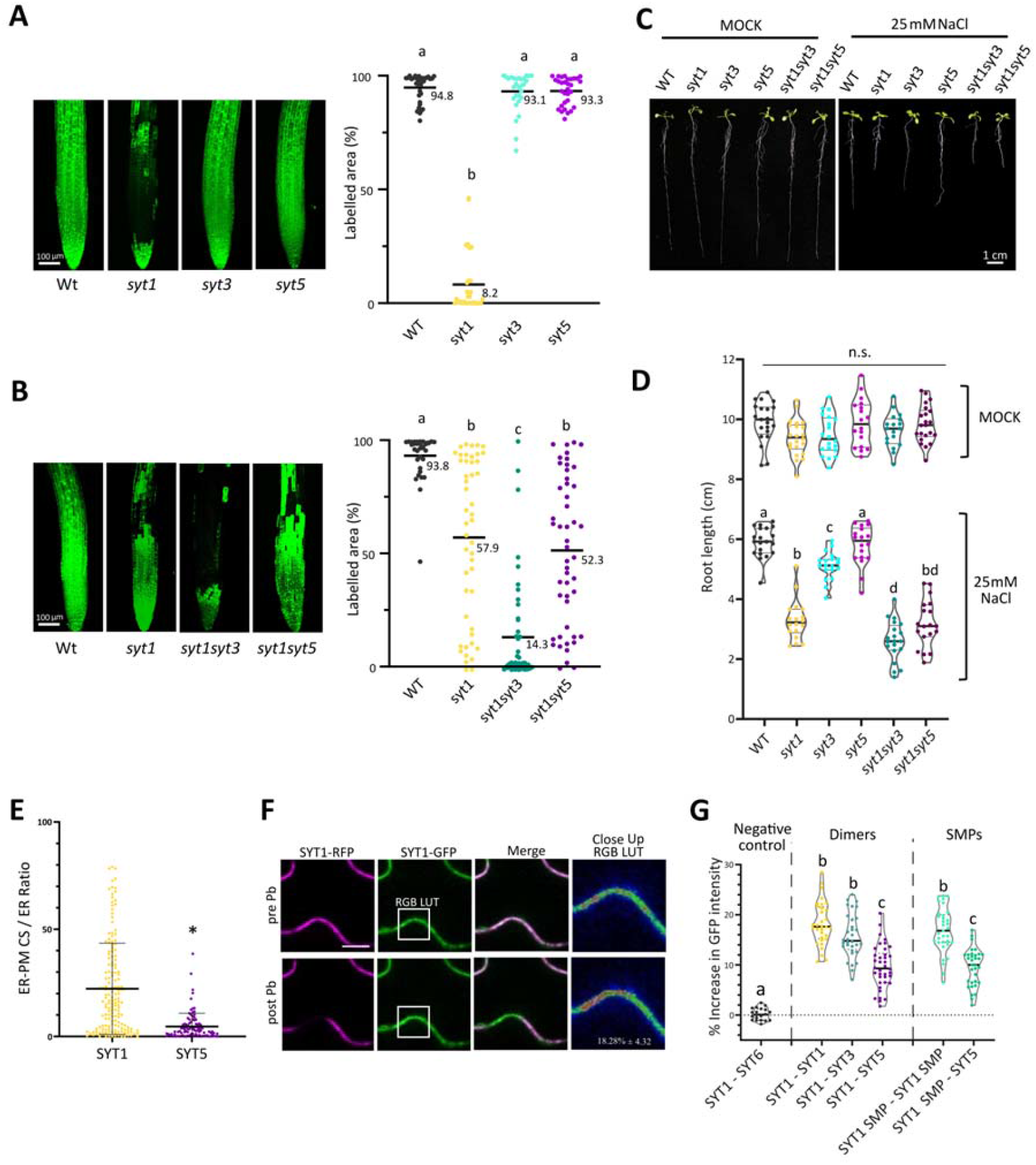
Mutations in *SYT1*, *SYT3*, and *SYT5* show different phenotypic outcomes to biotic and abiotic stress. A, B) Confocal images and cell viability quantification after 30min (C) and 10 min (D) of cold treatment in 6-d-old Arabidopsis roots. Seedlings grown in half-strength MS agar solidified medium under control conditions. Cell viability was determined by FDA staining and quantified as the percentage of root area with fluorescence above an automatic threshold stablished by the “Moments” algorithm of Fiji (see “Methods” for details). Scale bar = 100 μm. Each dot represents a measurement of an individual root. Horizontal lines represent mean values. Letters indicate statistically significant differences using one-way ANOVA Tukey multiple pairwise comparisons, p < 0.05 (n > 20). C). Root cell viability is compromised in the *syt1* mutant when treated at 8°C for 30 min, but not in the *syt3* or *syt5* mutants. *syt1syt3* mutant is more affected than *syt1* after 10 minutes at 8°C. *syt1syt5* has no additive phenotype compared to the single mutant *syt1*. C) 10 day-old seedlings grown vertically in control (left) and 25 mM NaCl (right) 1/10 MS plates under control conditions. D) Root length quantification from the WT and mutant seedlings grown in E). Letters indicate statistically significant differences using one-way ANOVA Tukey multiple pairwise comparisons, p < 0.05 (n > 20). E) ER-PM CS / ER ratio from segmentation was quantified in plants expressing SYT1 or SYT5. The asterisk indicates a statistically significant difference determined by a Student’s t-test, p < 0.05 (n > 20). F) FRET assay (Förster Resonance Energy Transfer) between SYT1-GFP and SYT1-RFP. Images pre- and post-photobleaching (Pb) were taken. Close Up corresponds with the box in SYT1-GFP and is represented by an RGB LUT to maximize the differences in GFP intensity. Scale bar = 10 μm. G) Förster resonance energy transfer (FRET) assays were conducted using SYT1, truncates SYT1-SMP, SYT3, SYT5, and SYT6 to determine the interaction between SYT1 and these proteins. Pairs of proteins were co-expressed in *N. benthamiana* leaves and analyzed at 2 dpi. RFP-tagged proteins (acceptor) were photobleached, and GFP-tagged proteins (donor) were quantified. The percentage increase in GFP intensity was calculated using the following formula: [(IAfter − IBefore) / IAfter] × 100, where IBefore and IAfter represent the means of the intensity from 6 measurements taken before and after photobleaching, respectively. The letters indicate statistically significant differences using one-way ANOVA Tukey multiple pairwise comparisons (p < 0.05).

The *syt1* mutant was identified by its hypersensitivity to grow at high NaCl (Schapire et al. 2008). Therefore, we examined the role of *SYT5* in NaCl tolerance by analyzing the root growth responses to NaCl treatments in the single *syt1, syt3,* and *syt5* and the double *syt1syt3* and *syt1syt5* mutants. In control conditions, without NaCl, no significative differences in root growth were found in any of the mutant genotypes compared with the WT (Fig. 1C, Fig. 1D). in the presence of NaCl, *syt1* and *syt3* mutants showed a strong and slight defect in root growth, respectively, while the growth of *syt5* resembled that of WT (Fig. 1C, Fig. 1D). Moreover, the *syt1syt3* double mutant but not the *syt1syt5* double showed an enhanced root growth defect compared to *syt1.* This suggests that *SYT1* and *SYT3*, but not SYT5, play redundant roles in cold and NaCl stress responses.

Loss-of-function *SYT5* mutants show increased disease susceptibility after dip inoculation with the Arabidopsis bacterial pathogen *P. syringae* pv tomato DC3000 (*Pst* DC3000) (S. Kim et al., 2021a, 2021b). Thus, to analyze whether *SYT1* or *SYT3* also play a role in resistance to *Pst* DC3000, we inoculated *syt1syt3* double mutant plants either by direct leaf infiltration or dip inoculation. At 4-dpi using either inoculation method, there was a similar growth of *Pst* DC3000 in *syt1syt3* as in wild type (WT) plants (Fig. S2A, Fig. S2B), indicating that, unlike *SYT5*, *SYT1* and *SYT3* do not play a role in plant resistance to *Pst* DC3000. These data support that *SYT1/SYT3* and *SYT5* play different roles in stress pathways in Arabidopsis.

The functional difference between SYT1/SYT3 and SYT5 could be explained, among other factors, by the proteins being localized in different membrane domains or forming different protein complexes. To address these questions, we used Arabidopsis stable transgenic lines expressing SYT1 and SYT5 fused to green fluorescent protein (GFP) under their endogenous promoters in their corresponding mutant backgrounds, *syt1* (Pérez-Sancho et al. 2015) and *syt5* (Lee et al., 2020). Confocal microscopy revealed that both SYT1 and SYT5 localize in the ER and ER-PM CS, although SYT1 exhibits a higher ER-PM CS / ER ratio than SYT5 (Fig. 1G, Fig. S3). Additionally, we used the SYT5:SYT5-GFP *syt5* to perform GFP-affinity purification followed by liquid chromatography-tandem mass-spectrometry (LC-MS/MS) analysis in order to identify the proteins physically associated with SYT5.

The results showed that SYT1 is one of the strongest *in vivo* SYT5 interactors (Table S1), which supports their localization in similar membrane domains. Such strong interaction, together with the identification of several other proteins known to be associated with ER/ER-PM CS (Table S1), suggests that both proteins are present in the same complexes. To investigate the in vivo direct interaction between SYT1 and SYT5, we conducted Förster Resonance Energy Transfer (FRET) assays. We generated *SYT1*, *SYT3*, and *SYT5* constructs with the *GFP* and *RFP* tagged at the C-terminus and transiently co-expressed them in *Nicotiana benthamiana* (Fig. S4). FRET analysis confirmed the *in vivo* interaction of full-length SYT1-GFP with SYT1-RFP (Fig. 1F, Fig. 1G), as well as with SYT3-GFP and SYT5-GFP (Fig. 1G). Because the E-Syts interact through their SMPs (Schauder et al., 2014), we investigated whether SYT1 and SYT5 could do so by using a SYT1 version lacking the C2 domains (SYT1-ΔC2AB-GFP). FRET analysis demonstrated the capacity of SYT1-ΔC2AB to both homo-oligomerize and to form hetero-oligomers with SYT5. These results strongly support that SYT1, SYT3, and SYT5 form complexes at the same ER-PM CS and that different localizations cannot account for the varied phenotypic responses observed in *syt1syt3* and *syt5*.

### SYTs of early land plants diverged into two evolutionarily conserved clades

The different functions exhibited by SYT1/SYT3 and SYT5 led us to analyze the sequences of the SYT family to identify potential divergences (Fig. S5A). This analysis revealed two clades: one including SYT1, SYT2, and SYT3, and a second clade comprising SYT4 and SYT5 (Fig. S5B), with the C2B domain showing the lowest identity between clades (Fig. S5C, Fig. S5D). Upon observing the divergence of SYTs in Arabidopsis, we investigated their phylogenetic relationship and their structural properties, exploiting the AlphaFold 2.0 predictions (Jumper et al., 2021). To identify SYT orthologs, we focused on those genes whose encoded proteins share a domain structure with those under investigation (Fig. S5A), including a predicted TM domain, an SMP lipid transport domain, and two C2 domains. A detailed phylogenetic analysis was performed as described in Methods. In addition to *Arabidopsis thaliana*, we used tomato (*Solanum lycopersicon*), the monocot *Brachypodium distachyon*, the liverwort *Marchantia polymorpha* and the green alga *Micromonas commoda* for the analysis (Fig. 2).

**Figure 2.**
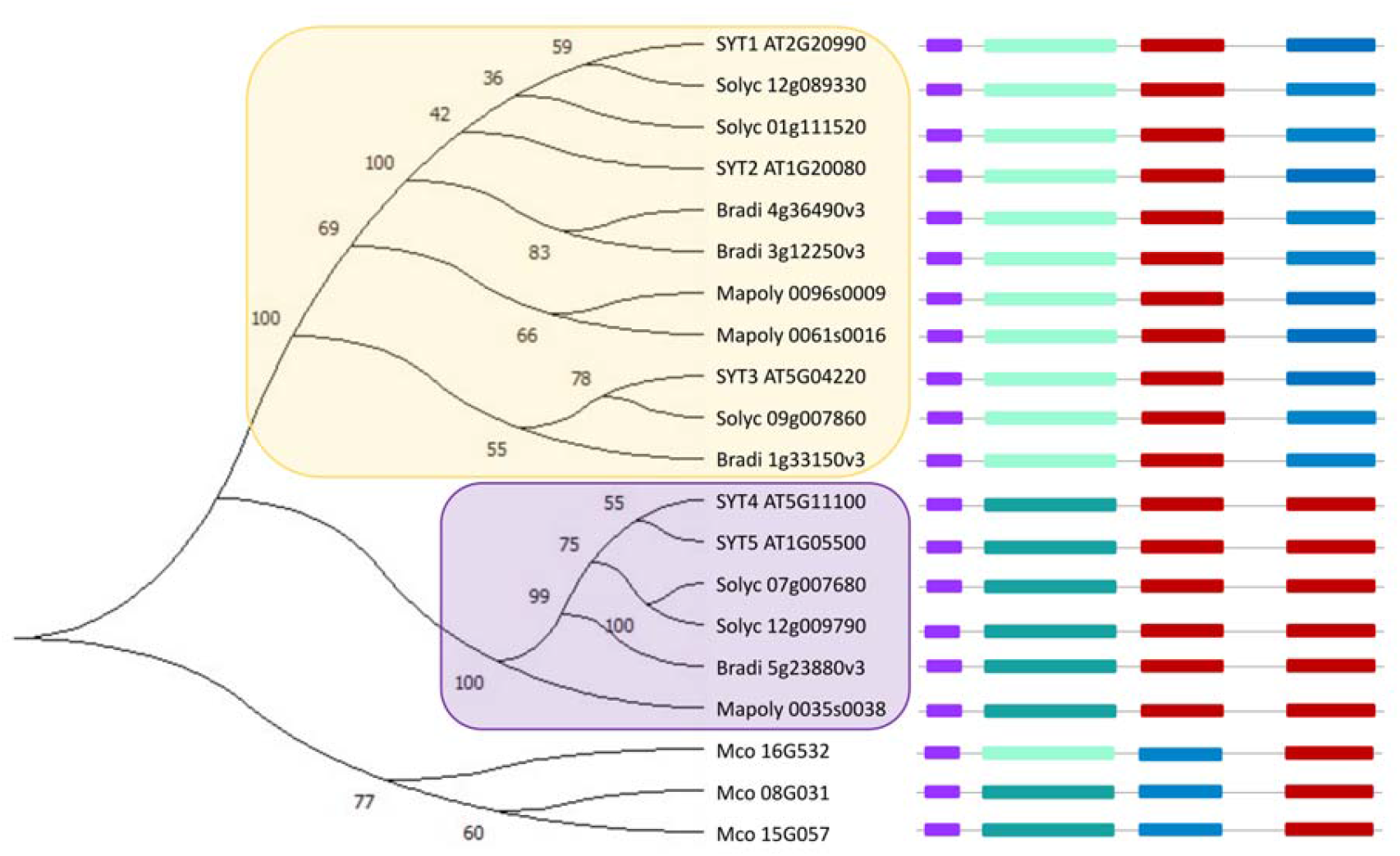
Phylogenetic tree of SYT1-like and SYT5-like proteins from the families of *Micromonas commoda*, *Marchantia polymorpha*, *Brachypodium distachyon*, *Arabidopsis thaliana* and *Solanum lycopersicum*. Protein structures featuring schematized domains are represented. The evolutionary history was inferred using the Maximum Likelihood method and the JTT matrix-based model. The percentage of replicate trees in which the associated taxa clustered together in the bootstrap test is displayed adjacent to the branches. SMP type of the SYT1-like clade is indicated in light green, while the SYT5-like SMP is in dark green. C2 domains binding Ca^2+^ are depicted in red, whereas those that do not bind calcium are shown in blue.

SYT homologs are already present in the green algae down to *M. commoda* that belongs to the Prasinophyceae (Chlorophyta), which are ancient members of the green lineage that diverged early from the lineage that led to all modern terrestrial plants (Qiu & Palmer, 1999). The three SYT-like proteins identified in *M. commoda* formed an outgroup of other plant species analyzed (Fig. 2). Interestingly, putative orthologs of SYT1 and SYT5 were identified in tomato, Brachypodium and Marchantia with their separation into two different clades. Monocots branched off from dicots 140-150 million years ago (Chaw et al., 2004) and tomato and *Arabidopsis* belong to two different families that diverged early in the radiation of dicotyledonous plants more than 90 million years ago (Ku et al., 2000). However, the finding of putative orthologs for SYT1 and SYT5 in Marchantia, the most basal lineage of extant land plants indicates that the speciation of the SYT family into SYT1-like and SYT5-like proteins occurred early in the evolution of land plants.

### Structural characteristics of SYT1-like versus SYT5-like clades

To investigate possible relationships between function and structure that characterize the SYT1-like and SYT5-like proteins, we compared the known structures of the SMP domain of human E-Syt2 (Schauder, Wu, Saheki, Narayanaswamy, Torta, Wenk, Camilli, et al., 2014) and the C2A domain of SYT1 (Benavente et al., 2021) with those predicted using AlphaFold (Jumper et al., 2021) for each member of the two groups. As expected, all the predicted proteins display the topology of those proteins determined experimentally: the SMP domains fold as an elongated dimeric structure that holds a hydrophobic tubular cavity to transport lipids (Fig. 3A left), and the C2 domains as a four-stranded beta-sandwich with a putative Ca^2+^-binding site placed in a cavity formed by three loops connecting the two beta-sheets (Fig. 3C left).

**Figure 3.**
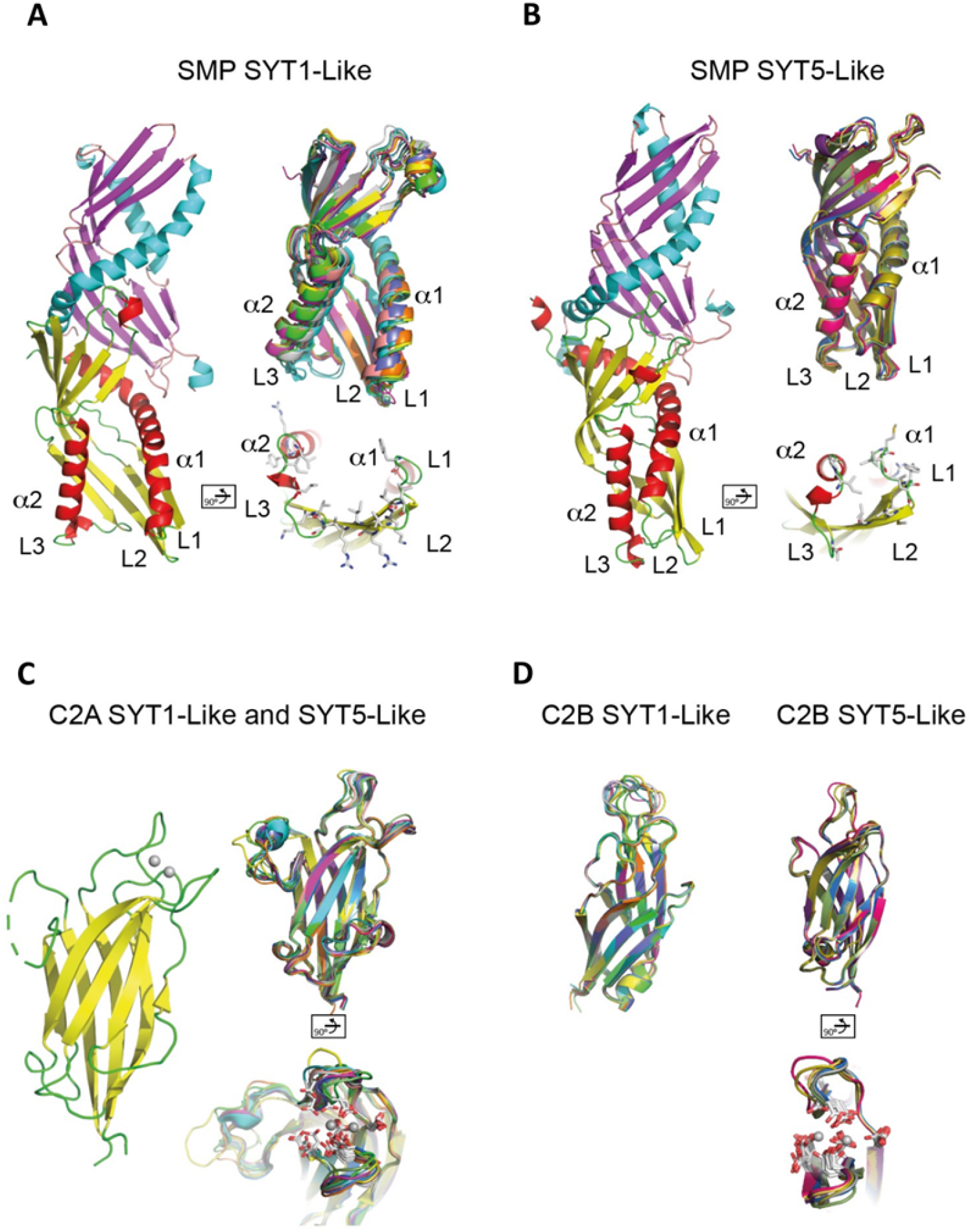
The predicted structures of SYT1-like and SYT5-like proteins. A) The structure of the SMP domain of SYT1-like (left) and SYT5-like (right) proteins. A view of the dimeric AtSYT1 and AtSYT5 together with the detailed structure of the “Tip” section highlights hydrophobic and positively charged residues and the superimposition of individual SMP domains of each group member. B) The crystal structure of the C2A domain of AtSYT1, left, and two views of the superimposition of individual C2A domains of each group member, right. The carboxylate side chains conforming the Ca^2+^ binding site are highlighted in a stick representation. Ca^2+^ molecules are represented by gray spheres. C) The superimposition of individual C2B domains of each member of the SYT1-like group, left, and two views of the members of the SYT5-like. The carboxylate side chains conforming the Ca^2+^ binding site are highlighted in a stick representation.

The structural alignment of the predicted SMP domains shows that proteins within each group exhibit a conserved structure, with the most significant variation present in the overall size and characteristics of the tip region of the SMP when comparing the two groups. These variations are primarily determined by the relative positions of α1 and α2 and by three loops (L1, L2, and L3) conforming the tip region (Fig. 3A, Fig. 3B, Fig. S6). This feature is relevant as the interaction of the SMP domain of E-Syts with the membrane occurs through the tip region, and the amino acid composition of these loops determines protein specificity for negatively charged PIPs (Wang et al., 2023). Aligned with this notion, the tip of the SMP of plant SYTs displays a pattern of solvent-exposed hydrophobic residues surrounded by polar residues (Fig. 3A, Fig. 3B, Fig. S6). This arrangement is well-suited for interactions with both the aliphatic components of the membrane and the charged polar heads of lipids. However, the polar residues in proteins of the SYT1-like group consist mainly of positively charged lysine residues, whereas neutral residues are present in proteins of the SYT5-like group. Therefore, the predicted structure and observed sequence variability in the tip region among proteins in each group suggest a unique membrane specificity for the SYT1-like and SYT5-like groups.

The C2A domain of SYT1 displays a canonical C2 structure with a well-defined Ca^2+^-binding pocket, which is involved in the binding of small and negatively charged phospholipids (Benavente et al., 2021; A. L. Schapire et al., 2008). In addition, it contains an unusually long loop adjacent to this site with an unknown function (Fig. 3C). The predicted structures and the analysis of the sequence alignment across all plant SYTs (i.e., SYT1 and SYT5-Like proteins) indicate the conservation of amino acids involved in Ca^2+^ binding (Benavente et al., 2021; Xu et al., 2014) and the presence of the long distinctive loop (Fig. 3C, Fig. S7). Therefore, the C2A domain of all plant SYTs is expected to display a common functional role. The structure of the C2B domain is predicted to show the folding of a C2 domain. However, while proteins in the SYT1-like group lack the essential residues for calcium binding, those in the SYT5-like group maintain them (Fig. 3D, Fig. S8). In this respect, it has been shown that the C2B domain of SYT1 displays constitutive and calcium-independent membrane binding properties (Benavente et al. 2021; Schapire et al. 2008), which could explain why SYT1 exhibits a more punctate pattern than SYT5 (Fig. 1G).

We additionally analyzed the predicted structures of extended synaptotagmins from *M. commoda*. Our findings revealed that they exhibit the same TM-SMP plus two C2 domains topology (https://www.uniprot.org/). However, in line with the phylogenetic analysis, the predicted structures differ from those predicted for proteins belonging to the SYT1-like or SYT5-like groups (Fig. S9). Interestingly, the C2A domains diverge from the conserved canonical C2 structure and lack the carboxylate side chains forming the Ca^2+^ binding site observed in plant SYTs. In contrast, the C2B domains are predicted to display a Ca^2+^ binding site.

Calcium signals function as specific regulators in diverse adaptation processes in plants (Tong et al., 2021; Tuteja & Mahajan, 2007). Consequently, proteins responsible for sensing these Ca^2+^ signatures have evolved to adjust their calcium-binding properties to decode such signals precisely (McAinsh & Hetherington, 1998). These unique calcium-binding properties and the underlying regulations of SYT1 and SYT5 may imply fine regulation at the subcellular level (Kudla et al., 2010; Steinhorst et al., 2022).

In this work, we show that the *syt1* and *syt5* mutants show different phenotypes supporting different biological roles for SYT1 and SYT5 proteins. Phylogenetic analyses revealed an early divergence of SYT1 and SYT5, and further structural studies predict distinctive properties between proteins present in SYT1 and SYT5 clades. These differences mainly rely on the Ca^2+^ binding characteristics of the C2B domains and the membrane binding moiety properties of their SMPs. Overall, our data suggest that the function specialization of the two SYTs clades might be a consequence of the diversity of land stresses that challenged early land plants (Delwiche & Cooper, 2015) which had to bear the specialization of plant mechanisms associated to membrane lipid homeostasis.

## METHODS

### Plant Material

*Arabidopsis thaliana* (Col-0 ecotype) and *Nicotiana benthamiana* served as the plant materials for this study. The Arabidopsis single mutant lines employed included *syt1-2* (AT2G20990) SAIL_775_A08 (previously characterized in Pérez-Sancho et al. 2015), *syt3-1* (AT5G04220) SALK_037933 (described in Ruiz-Lopez et al. 2021) and *syt5-2* (AT1G05500) SALK_036961 (described in Lee et al., 2020) obtained from The European Arabidopsis Stock Center (NASC: http://arabidopsis.info/). Double mutant *syt1syt3* was also previously reported in Ruiz-Lopez et al. 2021, while *syt1syt5* was obtained by crossing *syt1* and *syt5*. The identification of homozygous plants carrying the T-DNA insert in each mutant was confirmed through diagnostic PCR. Allele-specific primers, as listed in the supplementary primer table, were employed for this purpose. The transgenic lines used were SYT1:SYT1-GFP *syt1* (Pérez-Sancho et al. 2015) and SYT5:SYT5-GFP *syt5* (Lee et al., 2020).

### Plant manipulation and growth conditions

We followed established procedures for cultivating Arabidopsis. Seed sterilization was achieved by exposing them to a mixture of 100 ml commercial bleach and 3 ml 37% [w/w] HCl in a sealed container for 4 hours. After a minimum of 2 hours of air-clearing in a laminar flow cabinet, seeds were plated on 1/10 Murashige–Skoog medium. This medium included 1.5% (w/v) sucrose and 0.8% (w/v) physiological agar for solidification. A 3-day vernalization period at 4°C in darkness preceded the transition of plates to a vertical position in a growth chamber. This chamber maintained a long-day photoperiod (16 hours light / 8 hours darkness), a photon flux of 130 ± 30 mmol photons m–2 s–1, and a temperature of 22 ± 1°C. After 7 days of in vitro growth, seedlings were transplanted into pots containing an organic substrate and vermiculite [4:1 (v/v)]. These potted plants were maintained under controlled conditions (130 ± 30 mmol photons m–2 s–1 and a temperature of 22 ± 1°C) and watered every 2 days. For seed collection, dried seeds were stored in low-humidity conditions. Freshly harvested seeds were used for subsequent phenotypic analyses.

### Bacterial inoculation and disease resistance assays

Bacterial inoculations with *Pst* DC3000 were performed as previously described (Macho et al., 2007, 2010). Four- to five-week-old plants grown in short-day conditions were used. For direct leaf infiltration, rosette leaves were inoculated with a 5×10^4^ colony-forming unit (cfu/ml) bacterial suspension using a 1 ml syringe without needle. For dip inoculation, leaves were submerged for 30 seconds in a bacterial suspension containing 5×10^7^ cfu/ml and 0.025% silwet-L77. After inoculation (0 days-post inoculation, dpi) or 4 dpi, three 10-mm-diameter leaf discs were homogenized by mechanical disruption into 1 ml of 10 mM MgCl_2_. Then, bacteria were enumerated by plating serial dilutions onto LB agar. Bacterial enumeration was carried out in the dilution displaying between 50 and 500 colonies per plate.

### Cold treatments for cell viability in roots

Six-day-old seedlings of WT, *syt1*, *syt3*, *syt5*, *syt1syt3* and *syt1syt5* grown in control conditions were exposed to pre-chilled liquid 1/10 strength MS at 8°C for 10 minutes or 30 minutes. Following this, the seedlings were stained with 1/10 MS solution containing 10 μg/mL of FDA (Sigma, FDA stock solution 5 mg/mL in DMSO) for 5 minutes. Subsequently, the seedlings underwent a wash, and imaging was performed. In all instances, root imaging occurred within a 5-minute timeframe. The experiment was replicated three times with similar results.

### Salt stress root growth

Seeds were sown on 1/10 strength MS medium, 1.5% (w/v) sucrose, and 0.8% (w/v) agar and vernalized for 3 days at 4°C. After 2.5 days of growing in control conditions, seedlings were transferred to another control plate (Mock) or to a 1/10 MS medium with 25 mM NaCl (treatment) and kept at 22°C under a 16-h-light/8-h-dark cycle for another 8 days. Root length from at least 20 seedlings was measured. Three independent replicates showed consistent results.

### Plasmid constructs

Arabidopsis Col-0 served as the source for cDNA, acting as templates for the amplification of Coding DNA Sequence (CDS) regions of the target genes. High-fidelity DNA Polymerase (iProof from BioRad #1725301) was employed along with primers listed in Supplemental Table S2. pENTR L4-proCaMV35S-R1 (Karimi et al., 2007) and pENTR L4-UBQ10-R1 from (Alassimone et al., 2016), constituted the pENTR R4-promoter-L1 constructs. Simultaneously, the CDS without stop codons of SYT1 (1737 bp), its truncated version SYT1-SMP (711 bp), SYT3 (1620 bp), SYT5 (1680 bp) and SYT6 (2445 bp) were cloned into the pDONR L1-L2 vector, producing pENTR L1-CDS-L2 constructs. Validation of constructs was achieved through diagnostic PCR, restriction analysis, and sequencing. LR reactions (Invitrogen) were then used to combine plasmids pENTR L4-promoter-R1, pENTR L1-CDS-L2, and pENTR R2-tag STOP codon-L3 (RFP or GFP) with their corresponding pDestination (pDEST) vectors (pGWB5; pH7m34GW).

### Transient expression in *N. benthamiana*

Various constructs, including p19 (to counteract gene silencing associated with overexpression), were introduced into *Agrobacterium tumefaciens* (GV3101::pMP90) through electroporation. Subsequently, *A. tumefaciens* strains were employed for the transient transformation of *N. benthamiana*. The strains were cultured overnight at 28°C in Luria-Bertani (LB) medium supplemented with rifampicin (50 µg/ml), gentamicin (25 µg/ml), and the specific antibiotic corresponding to the construct (spectinomycin at 100 µg/ml or kanamycin at 50 µg/ml). Cultures were then centrifuged at 3,000 x g for 15 minutes at room temperature, and the resulting pellets were resuspended in an agroinfiltration solution (10 mM MES, pH 5.6, 10 mM MgCl2, and 1 mM acetosyringone). This suspension was incubated in the dark for 2 hours at room temperature. For experiments involving the expression of a single gene, the resuspended Agrobacterium cells were mixed to achieve an OD_600_ of 0.70 for the construct and 0.25 for the p19 strain. In double infiltration experiments, Agrobacterium strains were used at an OD_600_ of 0.40 for the constructs and 0.15 for the p19 strain. Two leaves from 3-week-old *N. benthamiana* plants (specifically, the 3^rd^ and 4^th^ leaves from the apex) were infiltrated on the abaxial side using a 1 ml syringe without a needle. The infiltrated plants were then maintained under growth conditions for 2 days before analysis by confocal microscopy.

### Confocal Microscopy Imaging

FDA, GFP or RFP fluorescence were detected using a Zeiss LSM880 confocal microscope. Excitation for FDA or GFP was achieved using the 488 nm argon laser, while the 561 nm laser was used for RFP excitation. Co-localization was assessed using sequential line scanning mode to separate signals. Fluorophore detection involved a PMT, a GaAsp (used to improve signal recognition), and an additional PMT for transmitted light. The objectives used were Plan-Apochromat 20X (water) and 40X (water) with up to a 4X digital zoom. Arabidopsis FDA-stained roots were visualized under the 20X lens. Z-stacks were taken starting from the surface of the roots and ending 25–30 μm deep inside the root (1 μm spacing). Localization of SYT1 and SYT5 in the transgenic lines SYT1:SYT1-GFP *syt1* and SYT5:SYT5-GFP *syt5* was observed in Arabidopsis cotyledons. For imaging on *N. benthamiana*, leaves were infiltrated as outlined in the “Transient Expression in *N. benthamiana*” section. 2 days post-infiltration, leaf disks were excised from the plants immediately prior to visualization, and lower epidermal cells were observed. All cortical plane images from Arabidopsis and Nicotiana represent a maximum Z projection of several planes from the cell surface to the interior. Equatorial images used in the FRET assay correspond to single-plane images. Microscopy image processing was conducted using the FIJI program (Schindelin et al., 2012).

### anti-GFP co-immunoprecipitation coupled with LC-MS/MS

Proteins interacting with SYT5-GFP were identified by anti-GFP co-immunoprecipitation (IP) followed by LC-MS/MS. 5–8 g of 10-day-old Arabidopsis seedlings stably expressing 35S::GFP (control) or SYT5::SYT5:GFP were used. Tissue was immediately frozen in liquid N2 after collection and frozen tissue was mixed with cold protein extraction buffer (50 mM Tris–HCl pH 7.5, 150 mM NaCl, 10% glycerol, 10 mM EDTA, 1 mM Na2MoO4, 1 mM NaF, 0.5 mM Na3VO4, 10 mM DTT, 0.5 mM phenylmethylsulfonyl fluoride (PMSF), 1% (v/v) protease inhibitor cocktail, and 1% NP-40) and incubated at 4 C for 45 min. Protein extracts were there incubated at 4 C for 1 h with GFP-Trap beads (Chromotek) and unspecific binding was minimized by several washes with wash buffer (the same as extraction buffer but with only 0.2 % NP-40). Bound proteins were eluted from the beads by incubating at 70 C for 20 min in 50 μl of SDS loading buffer, vortexing regularly. Immunoprecipitated proteins were separated on SDS–PAGE acrylamide gels, and each row of the gel was divided in 3 sections to maximize detection capacity: 1. above the band corresponding to SYT5-GFP, 2. around the band corresponding to SYT5-GFP, and 3. bellow the band corresponding to SYT5-GFP. Each of the three sections were digested and analyzed by tandem mass expectrometry independently.

### Arabidopsis eFP Browser Data Analysis

The expression levels of the five SYTs genes in Arabidopsis, spanning different tissues and developmental stages, were obtained from the accessible RNA-seq data on the eFP-Seq Browser website (https://bar.utoronto.ca/eFP-Seq_Browser/) (Sullivan et al., 2019). To assess the response to various abiotic stresses, 18-day-old wild-type (Col-0) seedlings were cultivated under long-day photoperiod, 24°C, and 50% humidity. The shoot outcomes of this experimental setup were retrieved from the Arabidopsis eFP Browser (http://bar.utoronto.ca/efp/cgi-bin/efpWeb.cgi) (Winter et al., 2007). Differential expression was determined by calculating the ratio of a gene’s expression value under a specific abiotic stress condition to its corresponding control value, thus providing the fold change in gene expression concerning the mock condition.

### In silico structural data analysis

Protein domain prediction was carried out utilizing the InterPro tool (https://www.ebi.ac.uk/interpro/) (Paysan-Lafosse et al., 2023). The structure of the proteins was calculated using the artificial intelligence algorithms provided by the AlphaFold server (Jumper et al., 2021). Sequence alignments were prepared using ESPript - https://espript.ibcp.fr (Robert & Gouet, 2014). The structural alignment of proteins and the cartoon figures were obtained using PyMOL (Schrödinger and DeLano 2020).

### Phylogenetic analysis

To unravel the evolutionary relationships among SYT1-like and SYT5-like proteins the sequences for all proteins were obtained from the UniProt database (https://www.uniprot.org/) (Consortium, 2023). Sequence alignment was carried out using the Clustal W tool (https://www.ebi.ac.uk/Tools/msa/clustalo/) (Sievers et al., 2011), and phylogenetic analyses were conducted by MEGA X software (Kumar et al., 2018). The evolutionary history was inferred using the Maximum Likelihood method and JTT matrix-based model. The bootstrap consensus tree inferred from 500 replicates represents the evolutionary history of the taxa analyzed. Branches corresponding to partitions reproduced in less than 50% bootstrap replicates are collapsed. The percentage of replicate trees in which the associated taxa clustered together in the bootstrap test (500 replicates) are shown next to the branches. Initial tree(s) for the heuristic search were obtained automatically by applying Neighbor-Join and BioNJ algorithms to a matrix of pairwise distances estimated using a JTT model and then selecting the topology with superior log likelihood value. This analysis involved 20 amino acid sequences. All positions with less than 95% site coverage were eliminated, i.e., fewer than 5% alignment gaps, missing data, and ambiguous bases were allowed at any position (partial deletion option). There was a total of 517 positions in the final dataset.

### FRET analysis

For Förster Resonance Energy Transfer (FRET) assays, 3-week-old *N. benthamiana* leaves were transiently co-infiltrated with proteins tagged at the C-terminal to GFP and RFP. Confocal images of single planes were obtained with the Zeiss LSM880 confocal microscope equipped with the Plan-Apochromat 40x/1.2 NA (water) objective lens. Equatorial sections of lower epidermal cells, where both proteins co-localized with comparable intensity values, were analyzed. Three distinct regions of interest (ROI) were defined for measurement purposes: ROI 1, where photobleaching was conducted over the acceptor fluorophore (RFP) using the 561 nm laser for 100 iterations at 100% power; ROI 2, a randomly selected area not photobleached; and ROI 3, designated as background with no signal. Within each ROI, six measurements of the donor fluorophore (GFP) were taken both before (Pre) and after (Post) photobleaching. The FRET efficiency was calculated as the percentage increase in donor fluorophore intensity (% ΔGFP) after acceptor removal, employing the formula: % ΔGFP = 100 x (Post-Pre)/Post (Liao et al., 2019). ROIs 2 and 3 served as technical controls, where no increase in GFP intensity was observed. More than 20 measurements were taken across different cells, leaves, and plants for each protein pair.

### SYT1 and SYT5 segmentation and ER-PM CS quantification

Confocal images of the cortical plane of SYT1:SYT1-GFP *syt1* (Pérez-Sancho et al. 2015) and SYT5:SYT5-GFP *syt5* (Lee et al., 2020) cotyledons of Arabidopsis plants were segmented in background, ER and ER-PM CS in a semiautomatic way, using the interactive machine learning tool ilastik (Sommer et al., 2011). ER and ER-PM CS areas were quantified in FIJI (Schindelin et al., 2012), and the ratio ER-PM CS / ER was calculated for each ROI.

### Statistic and graphs

Graphs and statistical analyses were carried out using Prism 8.02 for Microsoft (GraphPad Software, www.graphpad.com). The conducted tests included unpaired t-tests or one-way analysis of variance (ANOVA) followed by Tukey’s multiple comparison test (p < 0.05). In the figures, asterisks indicate statistical differences, as determined by the t-test (p < 0.05). For ANOVA analysis, distinct letters in the graphs signify statistically significant differences. The data represents the mean values; error bars illustrate the standard deviation (SD). All the experiments were replicated at least three times, yielding consistent results.

### Accession numbers

The genes investigated in this research are cataloged in The Arabidopsis Information Resource (https://www.arabidopsis.org/) with the corresponding accession numbers: *SYT1*: AT2G20990; *SYT2*: AT1G20080; *SYT3*: AT5G04220; *SYT4*: AT5G11100; *SYT5*: AT1G05500; *SYT6*: AT3G18370.

---

**Figure S1.**
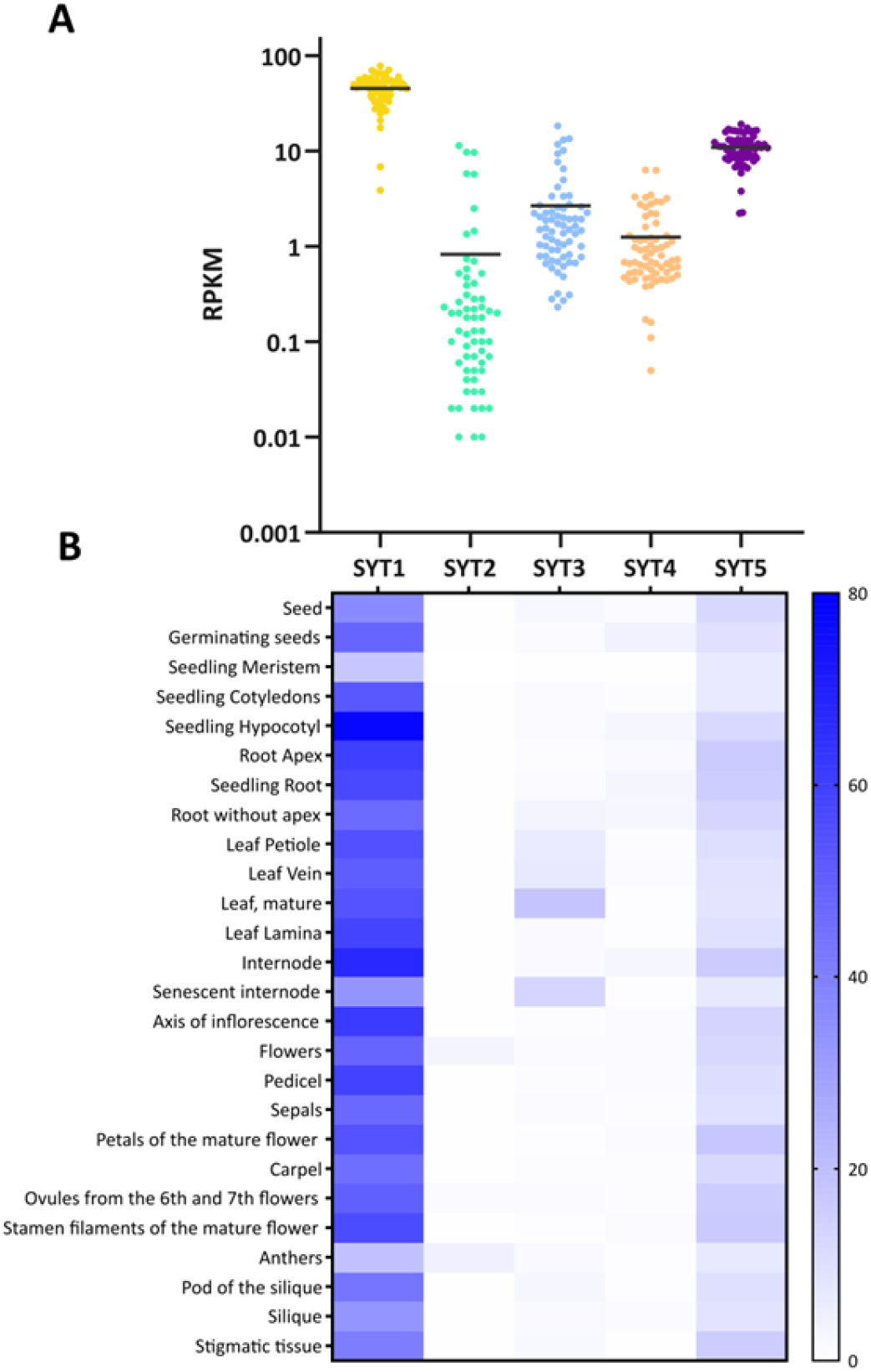
Expression levels of *SYT1, SYT2, SYT3, SYT4* and *SYT5*. A and B) RNAseq data of *SYT1, SYT2, SYT3, SYT4* and *SYT5* obtained from vegetative tissues at different developmental stages from eFP-seq Browser (https://bar.utoronto.ca/eFP-Seq_Browser/). A) Schematic summary of the level of expression of SYTs genes. Each dot represents a value of RPKM, and the bar represents the median, whose value is written above. B) Tissue expression represented by Heatmap.

**Figure S2.**
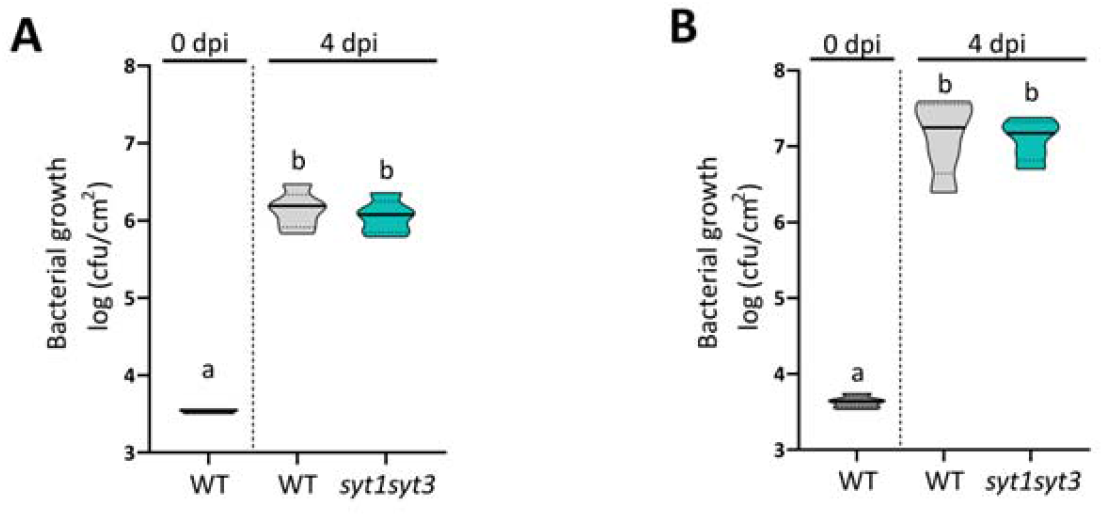
A, B) Growth of *Pseudomonas syringae* pv tomato DC3000 upon dip (A) or infiltration (B) inoculation in leaves of 4-to-5 week-old Arabidopsis plants at 0 or 4 days post-inoculation (dpi). Bacterial growth is represented as colony-forming units (cfu) per area of leaf tissue.

**Figure S3.**
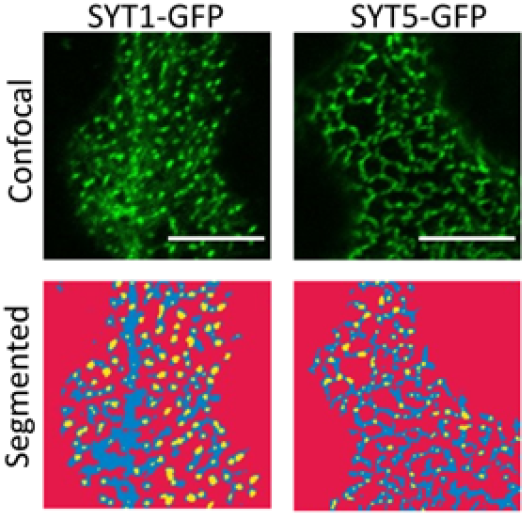
SYT1 and SYT5 are present at the ER-PM CS. Confocal images from SYT1-GFP *syt1* and SYT5-GFP *syt5* Arabidopsis epidermal cotyledon. Scale bar = 10 μm. The machine learning algorithm in the Ilastik tool calculates segmentation maps for Background (red), ER (blue), and ER-PM CS (yellow).

**Figure S4.**
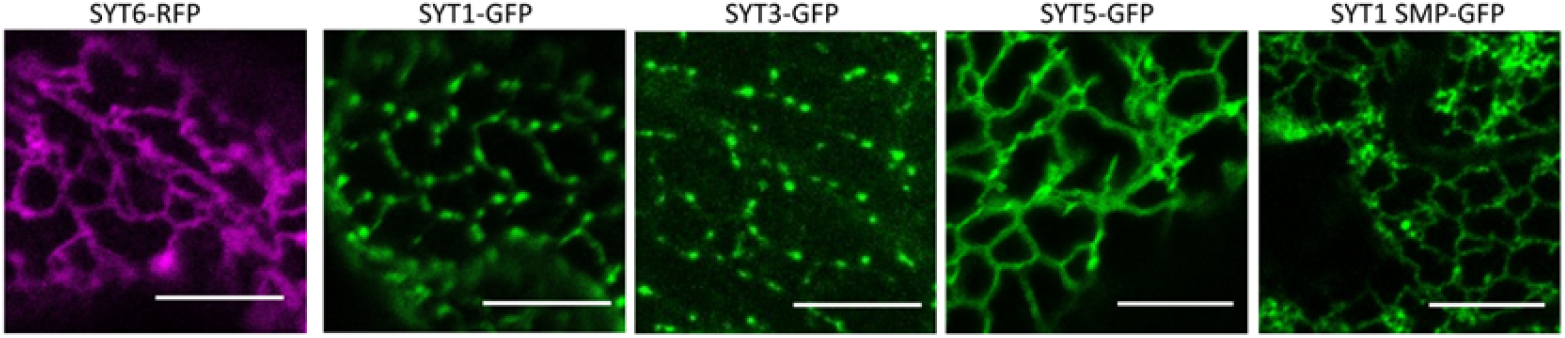
SYT3 and SYT5 interact with SYT1 through the SMP domain. Confocal images of 2 dpi *N. benthamiana* leaves expressing SYT6, SYT1, SYT3, SYT5, or SYT1 SMP, tagged to RFP or GFP at the C-terminal. The image corresponds to a Z-stack of the cortical zone of lower epidermal cells. Scale bar = 10 μm.

**Figure S5.**
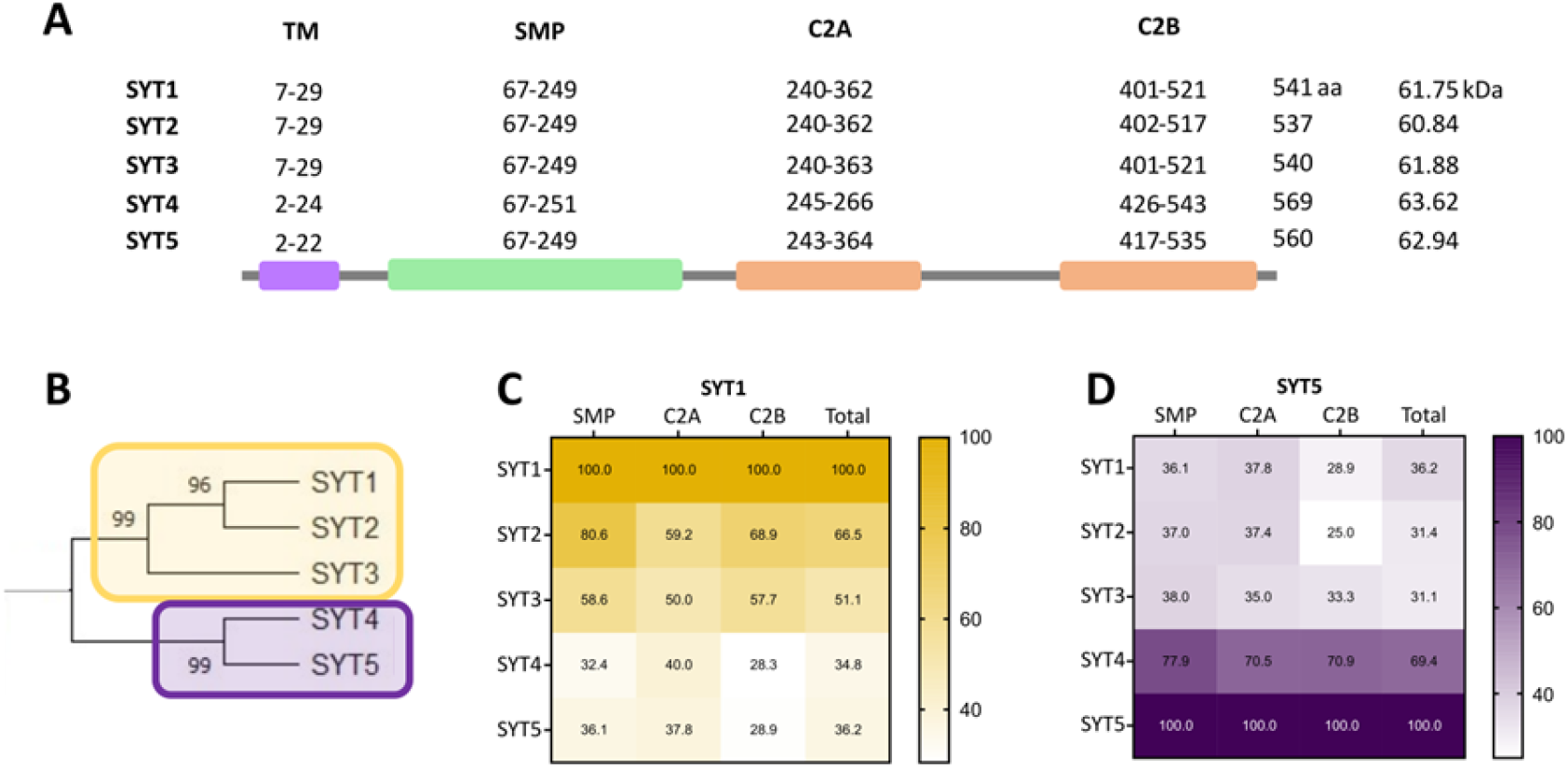
Arabidopsis SYT family protein analysis. A) Schematic view of SYT1-SYT5 proteins with domain data extracted from the UniProt database. The corresponding amino acid numbers indicate domain boundaries. B) Cladogram generated from the protein sequence alignment of Arabidopsis SYTs C, D) Protein sequence identity between SYT1 (C) or SYT5 (D) domains and the other SYT domains.

**Figure S6.**
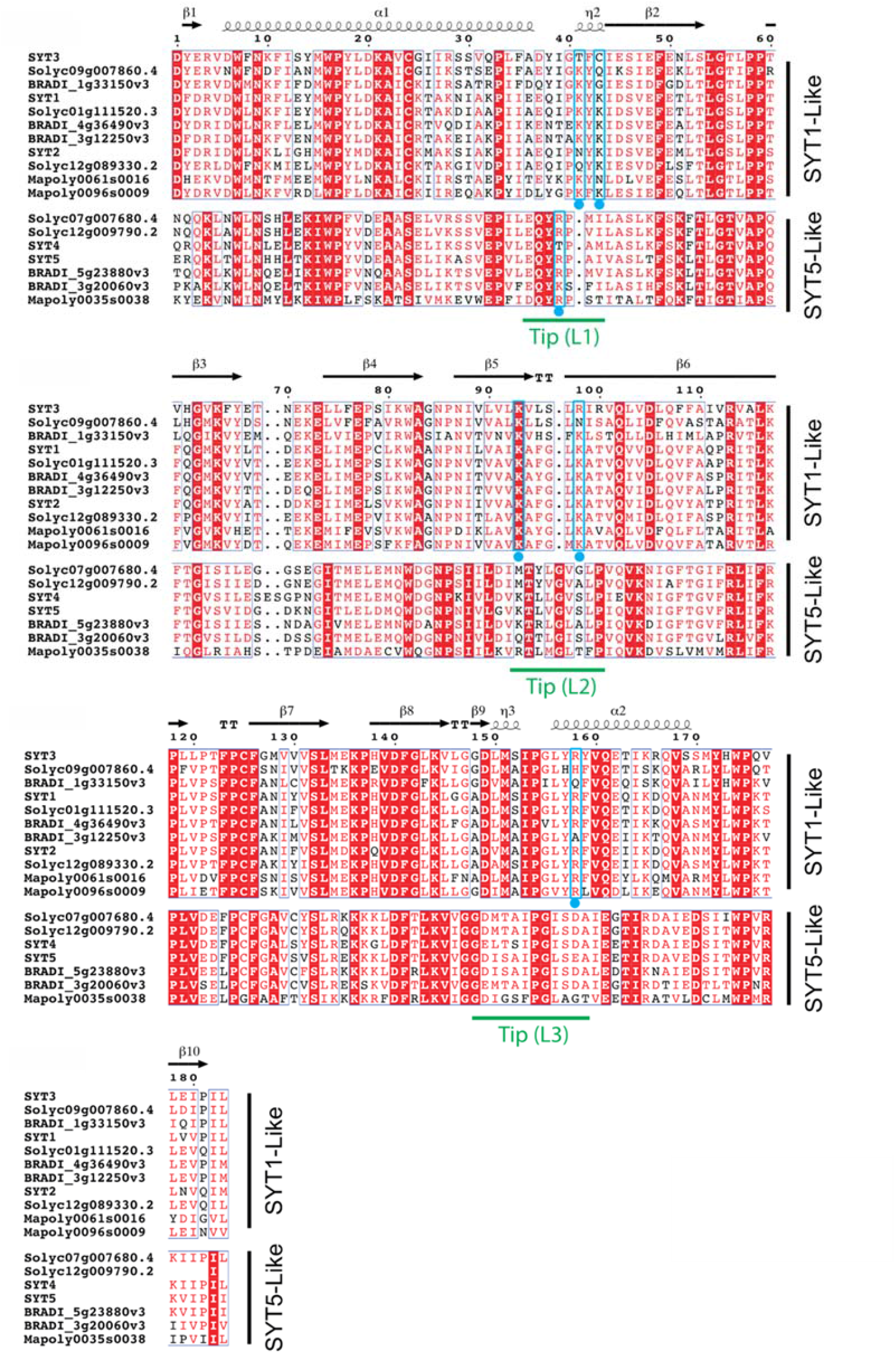
The Amino acid sequence alignment of the SMP, C2A, and C2B domains of plant SYTs. Residues involved in Ca^2+^ binding are indicated with a red sphere; residues likely to make up the basic sites are indicated with a blue sphere and highlighted in a blue box; secondary structural elements of the predicted structures are indicated. Root damage was quantified by assessing the percentage of the root area exhibiting visible FDA fluorescence above a predefined threshold, as previously described (Ruiz-Lopez et al., 2021).

**Figure S7.**
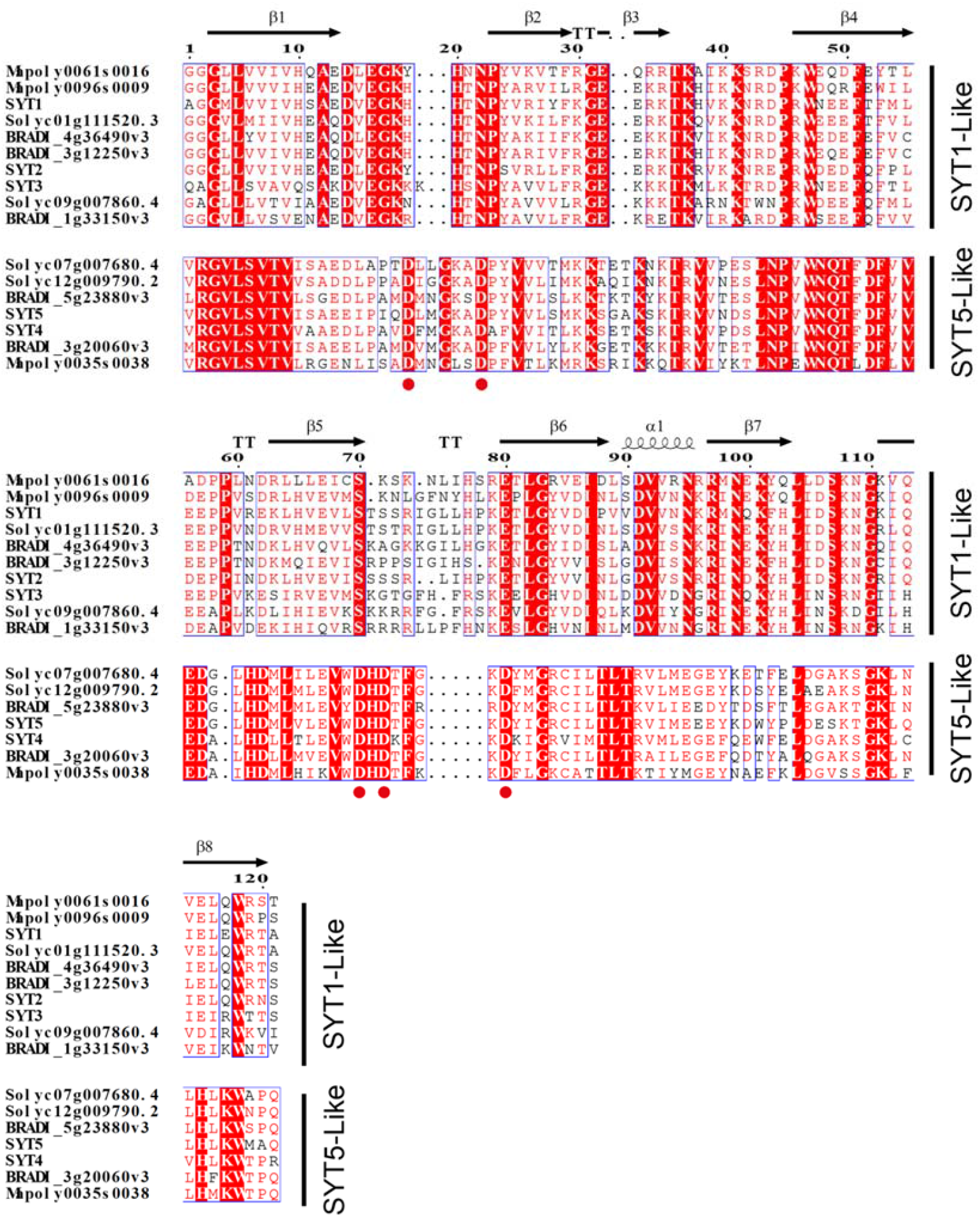
The Amino acid sequence alignment of the SMP, C2A, and C2B domains of plant SYTs. Residues involved in Ca^2+^ binding are indicated with a red sphere; residues likely to make up the basic sites are indicated with a blue sphere and highlighted in a blue box; secondary structural elements of the predicted structures are indicated. Root damage was quantified by assessing the percentage of the root area exhibiting visible FDA fluorescence above a predefined threshold, as previously described (Ruiz-Lopez et al., 2021).

**Figure S8.**
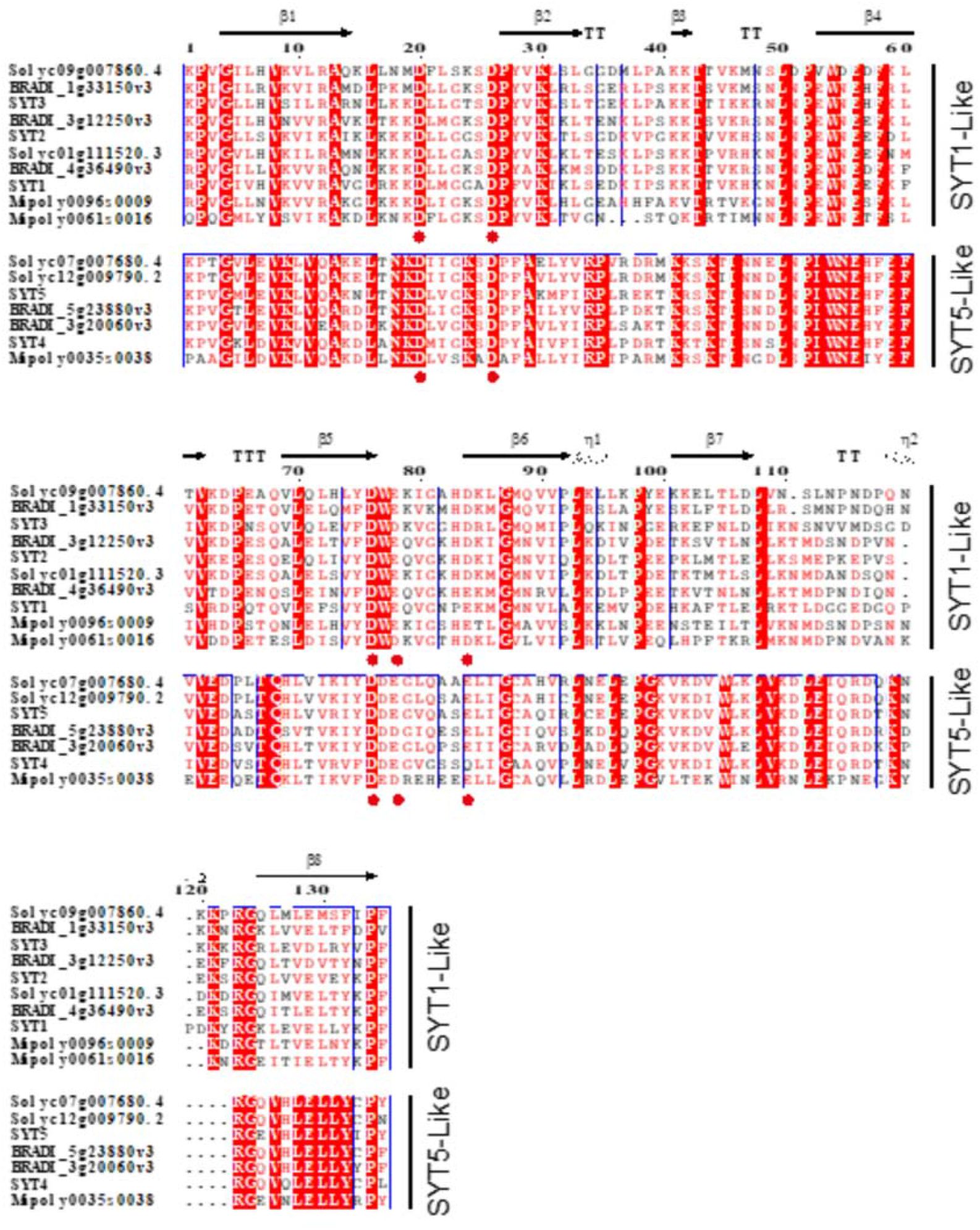
The Amino acid sequence alignment of the SMP, C2A, and C2B domains of plant SYTs. Residues involved in Ca^2+^ binding are indicated with a red sphere; residues likely to make up the basic sites are indicated with a blue sphere and highlighted in a blue box; secondary structural elements of the predicted structures are indicated. Root damage was quantified by assessing the percentage of the root area exhibiting visible FDA fluorescence above a predefined threshold, as previously described (Ruiz-Lopez et al., 2021).

**Figure S9.**
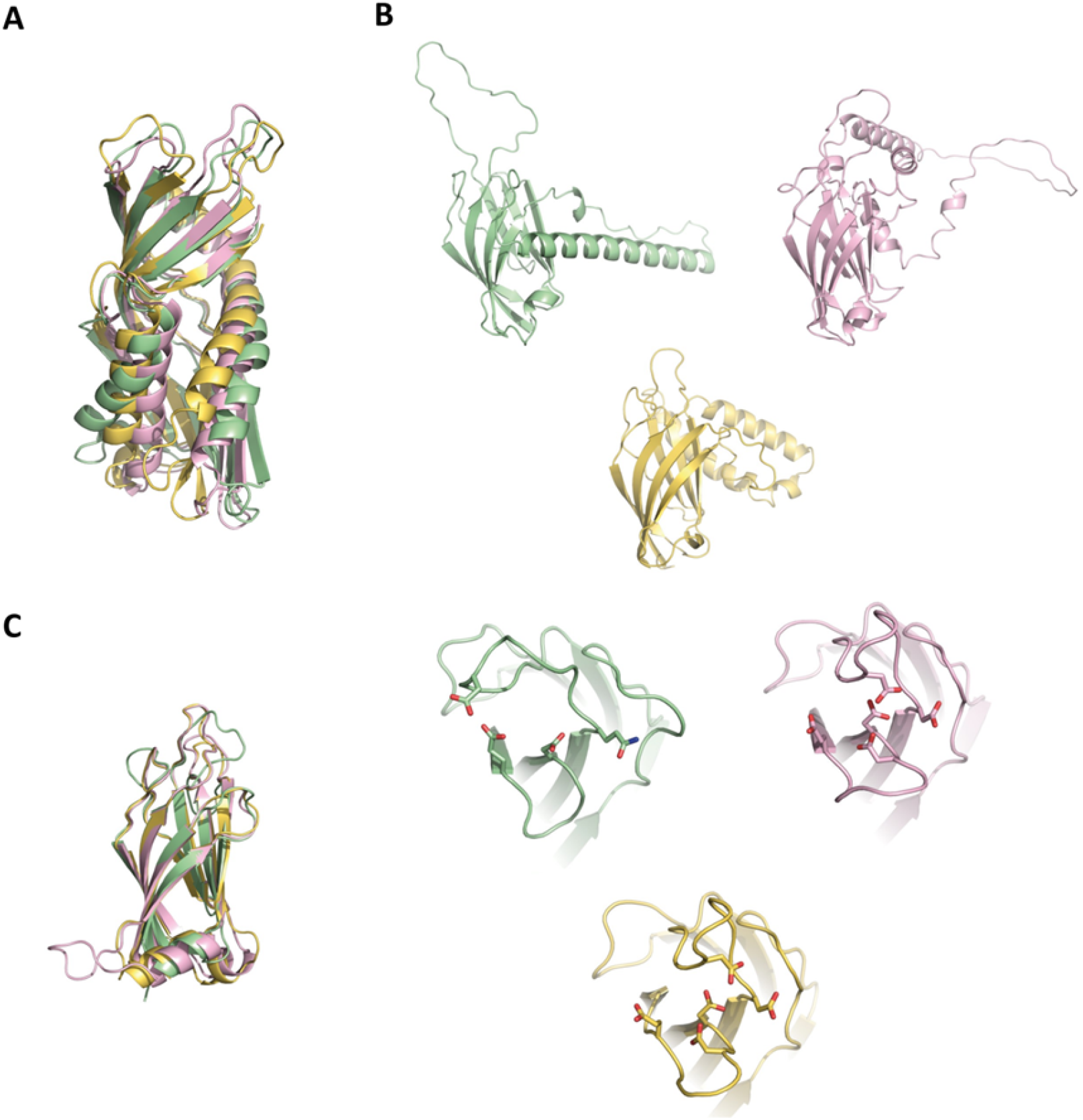
The predicted structures of the extended synaptotagmins of the green alga *Micromonas commoda*. Green (MCO 16G532) Yellow (MCO 08G031) and Pink (MCO 15G037). A) Superimposition of the SMP domains. B) A view of the C2A domains. C) Superimposition of the C2B domains (Left) and a view of the C2B domains showing the carboxylate side chains conforming the Ca2+ binding site (Right)

